# FlashRNA: An Efficient Model for Regulatory Genomics

**DOI:** 10.1101/2025.10.14.682350

**Authors:** Andrew J. Jung, Helen Zhu, Alice J. Gao, Roujia Li, Mykhaylo Slobodyanyuk, Vivian Chu, Declan Lim, Leo J. Lee, Albi Celaj, Brendan J. Frey

**Affiliations:** Deep Genomics; University of Toronto; Vector Institute

## Abstract

Transformer-based genomic sequence-to-function models effectively capture long-range genomic interactions but incur high computational costs due to the quadratic complexity of their self-attention layers. In this work, we introduce *FlashRNA*, which significantly improves computational and memory efficiency through *FlashAttention*, advancements in model architecture, and optimized training setup. *FlashRNA* achieves comparable or slightly improved predictive performance compared to similar sized *Borzoi* or *Flashzoi* models, notably without depending on pre-trained weights – a major limitation of *Flashzoi*. Remarkably, we trained *FlashRNA* from scratch in one day on a single GPU, significantly accelerating training and inference speed. These improvements can facilitate further developments in models for regulatory genomics by reducing computational cost. We demonstrate this in two downstream applications: 1) we train a large ensemble of 16 *FlashRNA* models and distill them into a single model to improve performance while maintaining efficiency, and 2) we fine-tune *FlashRNA* on three prediction tasks – ChIP-seq, RNA half-life, and translation efficiency – achieving performance matching or exceeding state-of-the-art task-specific models.

Code: https://github.com/deepgenomics/flashrna

## 1 Introduction

Recent transformer-based genomic models, like *Enformer* [2], *Borzoi* [14], *BigRNA* [4], and *AlphaGenome* [3], can accurately capture long-range cis-regulatory interactions across diverse cell states. Often classified as sequence-to-function models, they are trained to predict functional genome tracks from various experimental modalities measuring gene expression and epigenetic states, given an input genomic sequence. Unlike another class of genomic models that are trained on unlabeled sequences, sequence-to-function models can predict cell-state and disease specific effects of variants, along with their molecular mechanisms. Notably, they have shown potential as foundational models for DNA and RNA regulation, achieving state-of-the-art performance across a wide range of down-stream applications, from variant effect predictions, fine-tuning to capture cell-state-specific context of interest [13, 8, 19], and therapeutics design [4].

However, many promising applications of these genomic models rely on large-scale inference, such as interpreting variant effects genome-wide or designing nucleic acid sequences. One crucial challenge for these applications is the substantial computational cost, primarily due to the quadratic complexity of self-attention layers. Previous efforts to mitigate this issue with efficient self-attention mechanisms, such as FlashAttention [7, 6] or state-space models [9, 16], have resulted in either degraded performance on variant effect predictions[12] or dependency on pre-trained weights from existing models to work effectively [11].

In this work, we introduce *FlashRNA*, a novel approach to significantly improve computational and memory efficiency of genomic foundation models by leveraging *FlashAttention*, alongside improvements in model architecture and training strategy. Unlike prior approaches, *FlashRNA* does not depend on pre-trained weights [10], while preserving or improving downstream predictive performance. Remarkably, when matching parameters to *Borzoi*, we trained *FlashRNA* from scratch in just one day using a single Nvidia H100 GPU – a substantial improvement compared to 25 days with two A100 40GB GPUs for training Borzoi [14]. Furthermore, the reduction in computational requirements enables further improvements in scale and performance given an equivalent computational budget: we demonstrate this with an ensemble of 16 *FlashRNA* models and distillation of this ensemble into a single model that retains the ensemble’s predictive performance.

## 2 Methods

Our primary contribution is demonstrating that *FlashAttention* combined with Rotary Positional Encoding (RoPE) [17] can effectively replace computationally intensive self-attention layers of existing transformer-based sequence-to-function models without compromising performance or requiring pre-trained weights. Models like *Borzoi* and *Enformer* employ a hand-crafted relative positional encoding (PE), which incorporates domain-specific inductive biases such as decaying attention with distance. However, this custom positional encoding is incompatible with *FlashAttention*, necessitating an alternative. While RoPE is a broadly adopted relative PE compatible with *FlashAttention*, previous attempts resulted in inferior performance when training from scratch [10].

To compensate for losing the inductive biases from a domain-specific custom PE, *FlashRNA* employs aggressive data augmentation (shift margins up to 1,024 bases), higher dropout rates (0.3), and increased weight decay (0.1). Additionally, building upon the widely used U-Net based architecture used across many models, *FlashRNA* introduces the following improvements: GELU activation [10] replacing ReLU, Group Normalization [18] replacing Batch Normalization, and removing additive biases in linear layers.

In addition to improvements in the model architecture, we hypothesize that much of the information from CAGE and ChIP-seq tracks can be implicitly captured by RNA-seq, DNase-seq, and ATAC-seq alone. We trained *FlashRNA* exclusively on RNA-seq, DNase-seq, and ATAC-seq, which significantly reduced computational demands and improved convergence speed. The incorporation of *FlashAttention* also enabled substantially larger batch sizes and allowed the use of higher learning rates, further improving training efficiency and stability.

## 3 Results

We trained *FlashRNA* on a single Nvidia H100 GPU using the AdamW optimizer [15]. Compared to Borzoi, a similarly-sized FlashRNA model achieved 6.1x faster inference and 2.4x faster back-propagation, while handling 4.0x larger batch sizes and 3.5x longer sequences at inference (enabling context size up to 3.6 million base pairs), as shown in Figure 1. These speedups, in addition to other training improvements, enabled training *FlashRNA* from scratch in just one day on a single H100 GPU, compared to approximately 56 GPU-days required by Borzoi on an A100 40GB GPU.

**Figure 1.**
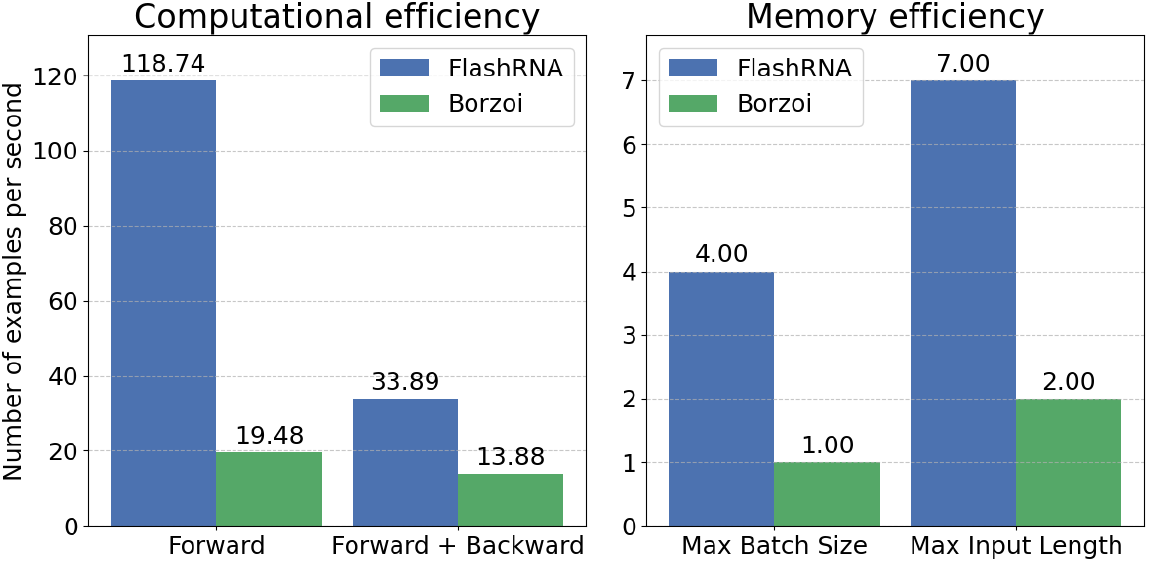
Computational and memory efficiencies of *FlashRNA* and *Borzoi*. (Left) examples per second during forward pass and combined forward-backward passes. (Right) the maximum batch size during training and maximum input length during inference relative to training context size before running out of memory. All measurements were made on a single Nvidia H100 GPU.

### 3.1 *FlashRNA*’s efficiency does not come at the cost of performance on key evaluation tasks

We evaluated *FlashRNA* on: 1) predicting coverage on held-out genomic intervals and 2) predicting variant effects on fine-mapped GTEx eQTLs [5]. On the task of predicting held-out test sets, *FlashRNA* demonstrated comparable predictive performance for RNA-seq, ATAC-seq, and DNase-seq tracks across four model replicates (Table 1). Similar results were also observed for predicting effect size on fine-mapped GTEx eQTLs, with *FlashRNA* and *Borzoi* achieving Spearman correlations of 0.406 versus 0.399, respectively (Table 2 right). Notably, excluding ChIP-seq and CAGE-seq tracks accelerated training without degrading performance on either held-out correlations or eQTL effect size predictions, suggesting these tracks provide minimal additional predictive value for these tasks. When predictions for these excluded tracks are needed, we demonstrate in Section 3.3 that pre-trained *FlashRNA* can be efficiently adapted through fine-tuning.

**Table 1.**
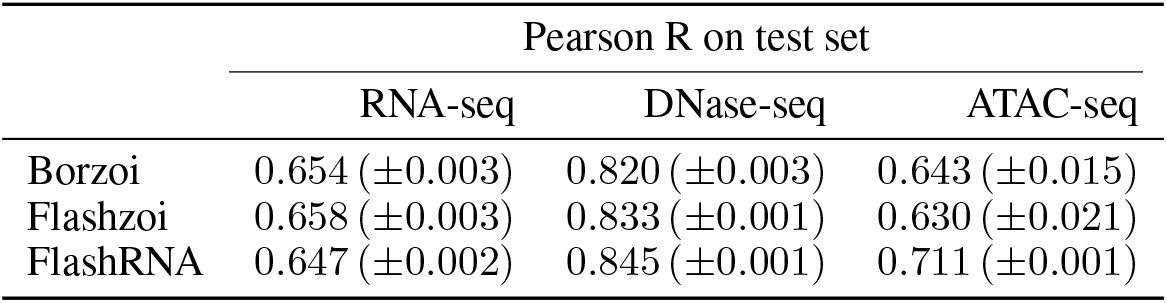
Predicting held-out tracks (the same ‘fold3’ test set used in *Borzoi*). Inverse normalization transformations, as described in [14], were applied to the model predictions and the target tracks before computing correlations. Pearson correlations for each of the four model replicates and their mean are shown.

**Table 2.** Predicting eQTL effect sizes. We used the same ‘logSED’ score used in *Borzoi*. Pearson correlations for each of the four model replicates and their mean are shown.

| | GTEx eQTL<br>Spearman $\rho$ |
| --- | --- |
| Borzoi | 0.399 ($\pm 0.003$) |
| Flashzoi | 0.398 ($\pm 0.012$) |
| FlashRNA | 0.406 ($\pm 0.005$) |

### 3.2 Distillation from a large ensemble improves performance while retaining efficiency

Leveraging *FlashRNA*’s computational efficiency, we trained a large ensemble of 16 models and investigated how performance scales with ensemble size. Model ensembling demonstrated substantial performance gains on the GTEx eQTL effect size prediction. Single *FlashRNA* models achieved a Spearman correlation of 0.406 while ensembles of 4 and 16 models achieved substantially higher correlations of 0.440 and 0.454, respectively – representing a 12% improvement from single to the full ensemble (Figure 2).

**Figure 2.**
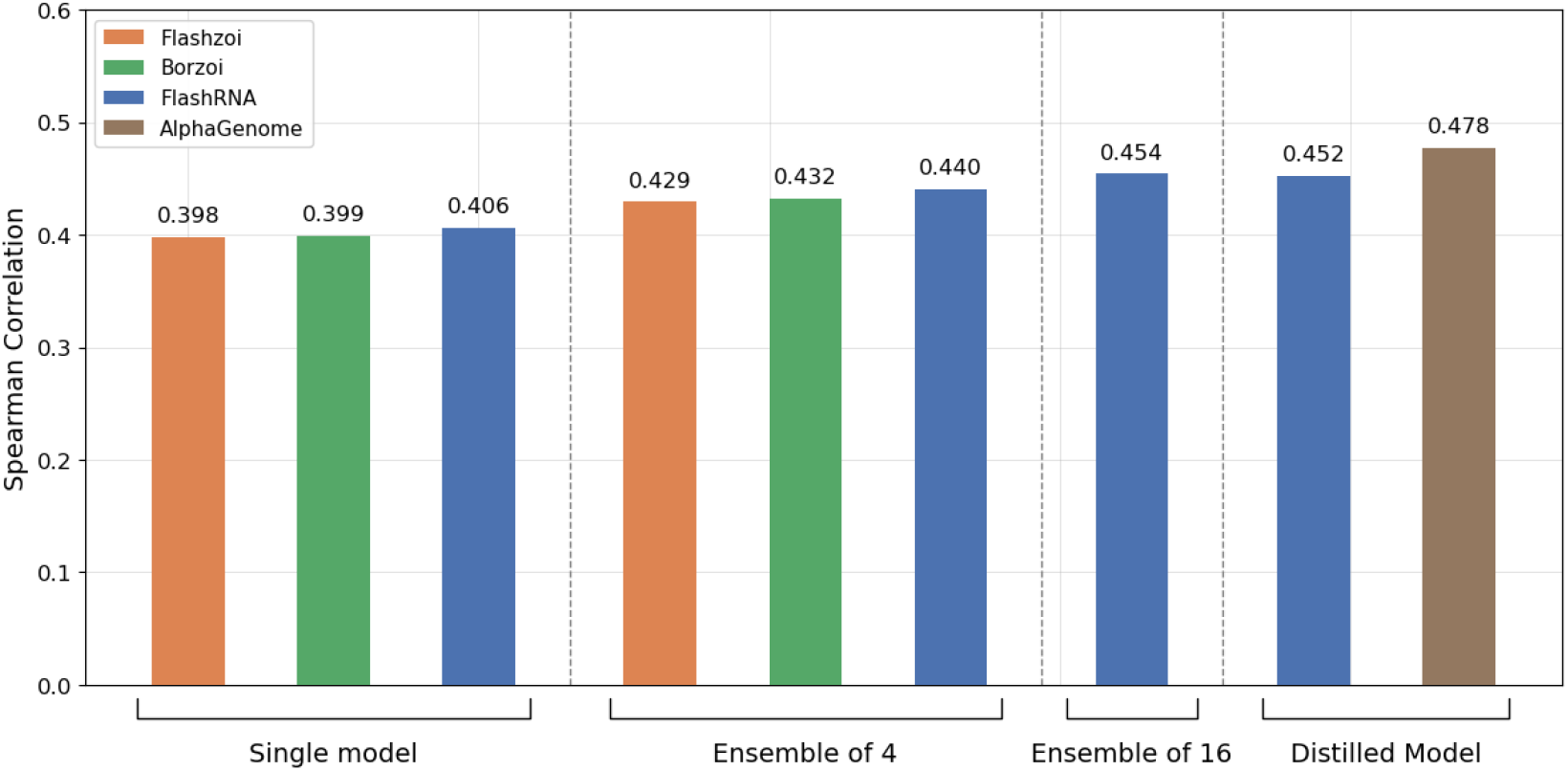
Spearman correlations of GTEx eQTL effect size predictions comparing *FlashRNA, Flashzoi, Borzoi*, and *AlphaGenome*. Single and an ensemble of four models are shown for *FlashRNA, Flashzoi, Borzoi*. For *FlashRNA*, we additionally trained an even large ensemble of 16 models and distilled it into a single model, retaining the ensemble’s performance. *AlphaGenome*’s distilled model is included for comparison but is only accessible via its API.

To maintain the performance benefit of ensembling while preserving computational efficiency of a single model, we distilled the 16-model ensemble into a single model. Results are summarized in Figure 2. The distilled model achieved a Spearman correlation of 0.452, matching the full ensemble’s performance while dramatically improving inference speed. This distillation approach significantly reduces the performance gap between *FlashRNA* and the current state-of-the-art model, *AlphaGenome*. Notably, *FlashRNA* achieves this competitive performance using less than half the parameters and training on standard GPU without sophisticated sequence parallelization – making it a more accessible option to researchers without specialized infrastructure.

### 3.4 *FlashRNA* enables efficient fine-tuning on downstream tasks

There has been significant interest in applying large genomic sequence-to-function models to a wide range of downstream tasks. While approaches like parameter-efficient fine-tuning have been proposed to reduce computational costs, full fine-tuning has been shown to achieve the best performance [19]. Here, we leverage *FlashRNA*’s efficiency to fine-tune it on three prediction tasks.

#### ChIP-seq

While *FlashRNA* was trained without ChIP-seq tracks for computational efficiency and faster convergence, we show it can be effectively adapted to predict transcription factor (TF) binding sites and histone modifications through fine-tuning. We add a ChIP-seq prediction head to the pre-trained *FlashRNA* and compare two fine-tuning approaches: (1) training only the prediction head while freezing the rest of the model, and (2) full fine-tuning.

After only 3 epochs of training, both approaches achieve competitive performance (Table 3). Full fine-tuning reaches near-parity with *Borzoi* (Pearson correlation of 0.593 vs. 0.595) while head-only fine-tuning achieves correlation of 0.574, consistent with the previous findings that full fine-tuning often achieves the best performance [19]. These results demonstrate that *FlashRNA*’s learned representations can be effectively adapted to predict new regulatory genomics tracks, despite their absence during pre-training.

**Table 3.** Fine-tuning performance on ChIP-seq tracks from the *Borzoi* dataset. Pearson correlations were computed on held-out tracks (the same ‘fold3’ test set used in *Borzoi*). Values show mean correlations and standard deviations across four model replicates.

|  | Borzoi | FlashRNA (head only) | FlashRNA (full) |
| --- | --- | --- | --- |
| Pearson R | 0.595 ($\pm 0.001$) | 0.574 ($\pm 0.001$) | 0.593 ($\pm 0.001$) |

#### RNA half-life and translation efficiency

Through pre-training on RNA-seq data, *FlashRNA* has likely learned latent representations relevant to various RNA properties. Here, we demonstrate that *FlashRNA* can be fine-tuned to predict RNA half-life and translation efficiency, achieving competitive performance with current state-of-the-art methods.

We fine-tune *FlashRNA* using the same datasets as *Saluki* [1] for RNA half-life prediction and *RiboNN* [20] for translation efficiency prediction, following their respective training and evaluation setups. Notably, while both *Saluki* and *RiboNN* rely on additional genomic annotations^1^ to boost performance, *FlashRNA* uses only RNA sequences. For each task, we add a task-specific head which takes pooling embeddings from *FlashRNA* as input.

As shown in Table 4, *FlashRNA* achieves performance comparable to both *Saluki* and *RiboNN*, despite using only sequences as input. Notably, *FlashRNA* significantly outperforms sequence-only versions of these models.

**Table 4.**
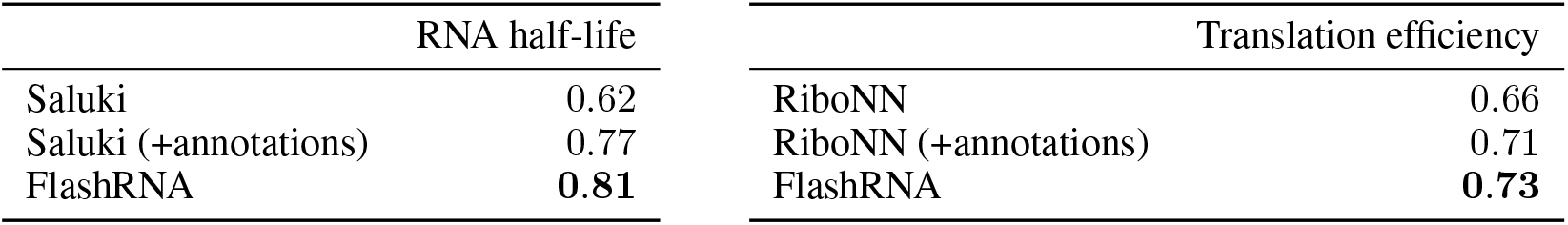
Performance on RNA property prediction tasks. Pearson correlation coefficients on human held-out test sets are reported. Models marked with ‘(+annotations)’ use sequence plus genomic annotations, while unmarked models use sequence only.

## 4 Discussion

In this work, we demonstrated how *FlashRNA* significantly enhances the computational efficiency of transformer-based sequence-to-function models. By leveraging *FlashAttention* and incorporating additional improvements in model architecture and training setup, *FlashRNA* achieves better performance on a fixed compute budget. Our results are consistent with recent findings in deep learning, where more general and computationally efficient architectures can replace specialized ones with domain-specific biases through data augmentation and regularization^2^.

The improved computational efficiency of *FlashRNA* enables many interesting future research directions, both for groups with limited computational resources and for those with greater resources who want to explore more broadly and iterate more quickly. These include exploring longer genomic contexts, improving resolution of model outputs, and applying the model to broader downstream tasks. By significantly reducing the computational barriers to training and inference, our work aims to facilitate further advancements in the field of modeling regulatory genomics.

## Footnotes

1 These include annotations for splice sites, codon reading frames, and UTR alignments.

2 For instance, recent gain in popularity of general transformer-based models where domain inductive bias is incorporated through data augmentation and training setup, compared to models with specialized equivariant architectures.

